# Non-affine displacements encode collective conformational fluctuations in proteins

**DOI:** 10.1101/840850

**Authors:** Dube Dheeraj Prakashchand, Navjeet Ahalawat, Satyabrata Bandyopadhyay, Surajit Sengupta, Jagannath Mondal

## Abstract

Identifying subtle conformational fluctuations underlying the dynamics of bio macro-molecules is crucial for resolving their free energy landscape. We show that a collective variable, originally proposed for crystalline solids, is able to filter out essential macro-molecular motions more efficiently than other approaches. While homogenous or ‘affine’ deformations of the biopolymer are trivial, biopolymer conformations are complicated by the occurrence of in-homogenous or ‘non-affine’ displacements of atoms relative to their positions in the native structure. We show that these displacements encode functionally relevant conformations of macromolecule and, in combination with a formalism based upon time-structured independent component analysis, quantitatively resolve the free energy landscape of a number of macromolecules of hierarchical complexity. The kinetics of conformational transitions among the basins can now be mapped within the framework of a Markov state model. The non-affine modes, obtained by projecting out homogenous fluctuations from the local displacements, are found to be responsible for local structural changes required for transitioning between pairs of macro states.

## Introduction

The conformational landscape underlying the dynamics of a bio macromolecule is complex and generally composed of rough terrains and shallow valleys. Resolving key basins corresponding to the metastable conformational states of the biomacromolecule in the landscape is a nontrivial task and has remained a topic of contention over the last few decades. A common approach in this direction has been the projection of the biomolecular phase space along judiciously constructed collective variables (CV). A clever choice of CV makes it possible to explore the mechanisms underlying conformational changes. A good CV^1^ is one that a) provides dimensionality reduction b) based on dynamics and c) can predict future events. Traditionally intuition-based CVs have not only catered to the needs of deciphering the underlysing pathways but also helped in efficiently biasing the MD simulation in enhanced samplings like umbrella sampling and metadynamics, and also building Markov state models (MSM).^2–10^ However, the heterogeneity associated with individual bio macromolecule has made the choice of CV non-unique, for optimally describing biomolecular conformation. This has resulted in the proposition of diverse sets of CVs over the years for capturing the biomolecular conformational dynamics. Many methodological developments have helped in devising an optimal CV for numerous complex systems. Spectral gap optimization of order parameter (SGOOP)^6^ algorithm proposed by Tiwary and Berne helps in capturing the combination of CVs that record the slowest dynamics of the system. Using a combination of time-structured independent component analysis (TICA) and metadynamics^5^ approach it has been shown that it is plausible to sample the slowest events and additionally drive the system along the slowest modes for enhanced sampling. Parrinello and coworkers, ^3^ have recently developed an approach where initially a metadynamics is carried out using a combination of commonly used CVs and subsequently a variational TICA approach on this sampling reveals slow modes involved in the system. Then futher metadynamics is performed using these slow modes as CVs. However, the underlying complexity of the conformational landscape has also necessitated the emergence of machine learning based approaches^11–14^ for CV optimisations.

On the other hand, in a crystalline solid, there exists an attractive prescription ^15–18^ to resolve its overall displacement from a reference lattice into two displacement subspaces: a) “affine” displacement, which is the trivial homogenous deformations from the reference lattice and b) “non-affine” displacement, which is the remainder of the deformations (see Figure 1). This formalism reveals that the necessary precursors for the initiation of crystal lattice defects such as dislocation dipoles which are provided by the thermally excited non-affine fluctuations.^16^ Application of this formalism revealed that rigid solids ^19^ are thermodynamically metastable even at infinitesimal stress, thereby enabling us to conclude that non-affine fluctuations are crucial in releasing the stress exerted on the yielding crystal. This helps rigid solids under external loading to decay from the metastable state to a stable state through a first order phase-transition. Using the non-affine formalism it is also possible to describe the non-trivial singular deformations like pleats and ripplocations^20,21^ undergone by solids having atoms bound by strong chemical bonds, which disallow any kind of dislocations. Also it has been shown, ^18,22^ that colloids can be ordered in any given crystal structure by suppressing the non-affine displacements, with the help of special feedback-control laser traps. Using a similar projection formalism as described in Ganguly et al., ^15^ for large bio macrolecules lacking any kind of long-range structural order it has been shown by Dube et al., ^23^ that important conformational changes are always accompanied by non-affine fluctuations and that regions in proteins with high susceptibility for non-affine fluctuations correlate with binding hotspots and the magnitude of the spatial correlations reveal allosteric connections in different locations of protein molecules.

**Figure 1:**
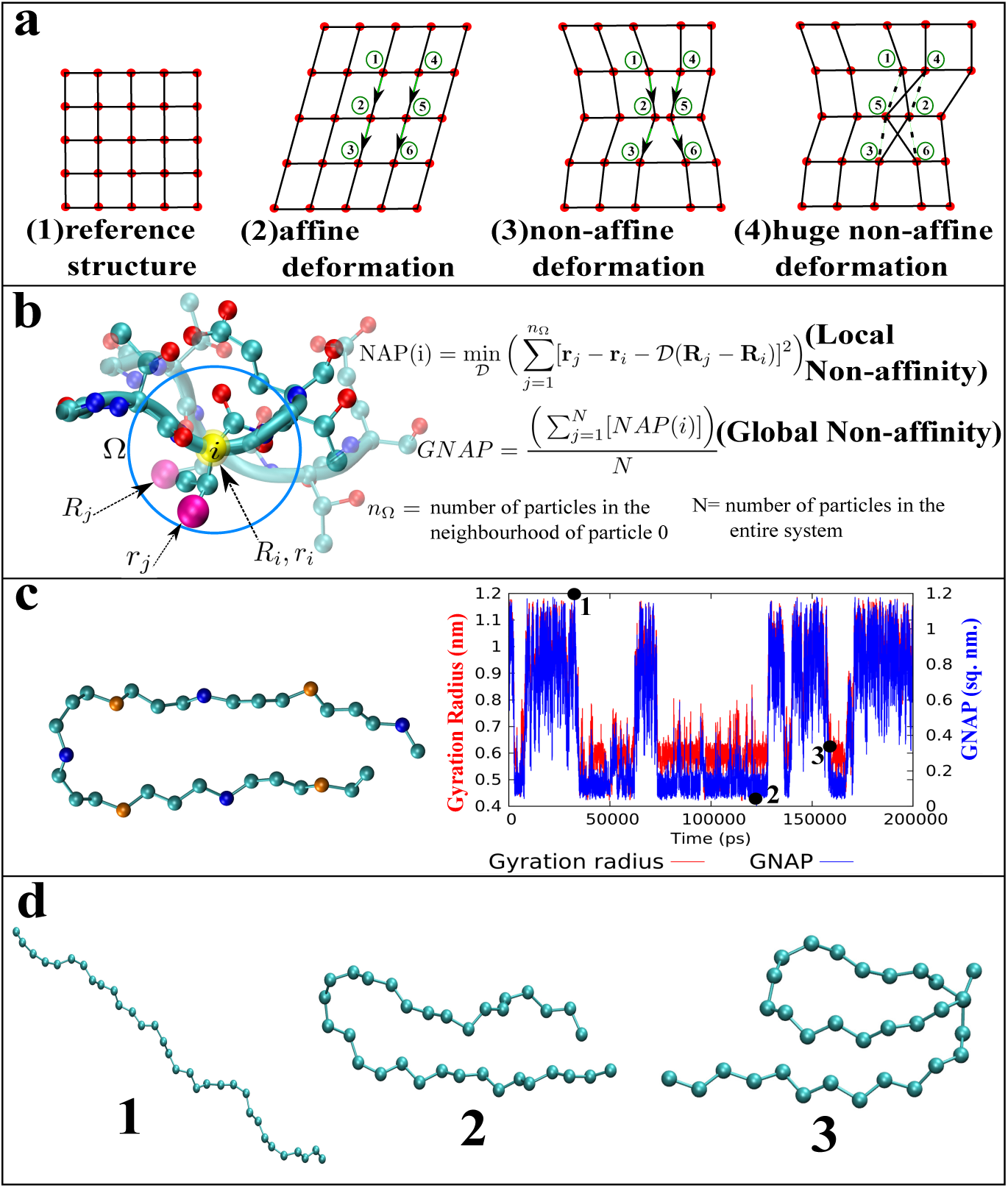
**a.** Illustration of ‘affine’ and ‘non-affine’ deformation in a lattice. **b**. Extension of the concept of non-affinity to protein **c**. The time profile of radius of gyration and GNAP (relative to beta-hairpin conformation) of a 32 bead charged-neutral polymer. The blue beads are positively charged beads while the orange ones are negatively charged. **d**. The representative snapshots corresponding to different GNAPs in the part c

As will be detailed below, the current work transitions this idea of ‘non affine displacement’, originally proposed for crystalline solids, to bio-macromolecules. We show that collective variables based upon non affine displacements in a bio-macromolecule, called ‘NAP’ and ‘GNAP’, can capture the crucial structural transitions in bio-macromolecules. In combination with time structured independent component analysis (TICA)^24,25^ and Markov-state model, GNAP provides a theoretically robust strategy for identifying key conformational motion in protein.

## Theory

Non-affine displacements in a bio-macromolecule are derived by filtering out routine homogenous displacements (known as ‘affine deformation’) from reference structure. Figure 1a-b schematically introduces us with non-affine displacements as well as the local and global non-affine parameter^15–17^ (NAP and GNAP respectively). Figure 1a illustrates affine and non-affine deformation for a crystal. Figure 1b illustrates the formalism in a protein: ^23^ Consider a neighbourhood Ω_*i*_ centered around an atom *i*. The instantaneous positions of the *j* = 1, *n*_Ω_ atoms in Ω_*i*_ may be compared with the set of fixed reference positions **R**_*i*_. The reference atomic positions, for example, could represent the native state of the protein. NAP defines the least square error made by trying to define **r**_*i*_ as a “best fit” affine deformation 𝒟 of **R**_*i*_ viz. 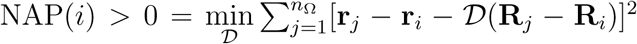. Here the minimisation is over choices of the deformation matrix 𝒟.^15^ Thus *NAP* (*i*) quantifies the “local structural changes” of i-th particle of the biomolecule. The central quantity of our current work, the global non-affine parameter (GNAP), is the average of NAP over the macromolecule (Figure 1b): 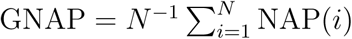

One can show^15,17^ that the thermal average of the local non-affine parameter is given by the trace of the displacement co-variance matrix projected onto the non-affine subspace viz. ⟨NAP(*i*)⟩ = Tr P[C(*i*)]P. Here C(*i*) = ⟨Δ^T^(*i*)Δ(*i*)⟩ is the displacement covariance with Δ(*i*) consisting of the relative displacements **u**_*i*_ − **u**_*j*_ of the atoms from the reference positions arranged as a column vector with 3*n*_Ω_ elements. The projection operator P depends *only* on the reference configuration and is obtained by first defining the 3*n*_Ω_ × 9 dimensional block matrix R with elements R_*jα,γγ*′_ = *δ*_*αγ*_(*R*_*jγ*′_ − *R*_*iγ*′_), where the Greek indices correspond to the spatial dimensions and take values 1, 2 or 3. Finally, P = P^2^ = 1 − R(R^T^R)^*−*1^R^T^.

While the sum of the eigenvalues of P[C(*i*)]P determines the thermal average of NAP, the corresponding *eigenvectors* govern the atomic conformations which lead to non-affine displacements. For example, in crystals, these non-affine modes with the largest eigenvalues correspond to precursors to crystal defects^16,17^ and are implicated in plastic events and loss of rigidity.^26^ In proteins too, they correspond to local conformational changes which are important for ligand binding and allostery. ^23^ One may also characterize regions in the protein which are particularly amenable to non-affine displacements using the local NAP susceptibility defined as ⟨NAP(*i*) −⟨NAP(*i*)⟩)^2^⟩. In this sense, the NAP susceptibility behaves as any other thermodynamic local response function such as heat capacity or compressibility. We have implemented the computation of GNAP for any macromolecules in popular MD analysis tool PLUMED2. ^27,28^

## Simulation method

We have used computational models of three different biomacromolecules in our current work: a) 32-bead model polymer with partial positive and negative charges along different beads, b) C-terminal domain of GB1 *β*-hairpin (residues 41–56) and c) Bovine pancreatic trypsin inhibitor (BPTI) protein. Two of these systems (polymer and *β*-hairpin) have been the subject of prior computational studies ^7,29^ by our group. (see Figure 2) In the current work, we have employed same simulation protocols as have been done earlier. The polymer mimics bead-in-a-spring model with a total of 32 beads connected by harmonic bonds and angles, the details of which are documented else-where. ^29^ The polymer consists of alternative positive and negative charges on the beads in a periodic fashion. Apart from the electrostatic interaction, the polymer beads additionally interact with each other and the aqueous media via lenard Jones (LJ) interaction. The LJ parameters of the polymer beads have been kept fixed at *σ* = 0.4 nm and *ϵ*_*p*_ = 1.0 kJ/mol. The other cross polymer-solution LJ parameters have been determined by geometric combination rule. As in previous case, the solvent has been modelled as SPC/E water model.^30^ All simulations have been done with a single polymer chain inside a cubic box of volume ≈ 125 nm^3^ consisting of 3850 water molecules at an average temperature of 300 K and an average pressure of 1 bar using NPT ensemble. The details of the thermostat and barostat in addition to other simulation details can be found in our previous work.^29^ We have run 200 independent simulations, each 30 ns long, by starting with diverse conformation of the polymer.

**Figure 2:**
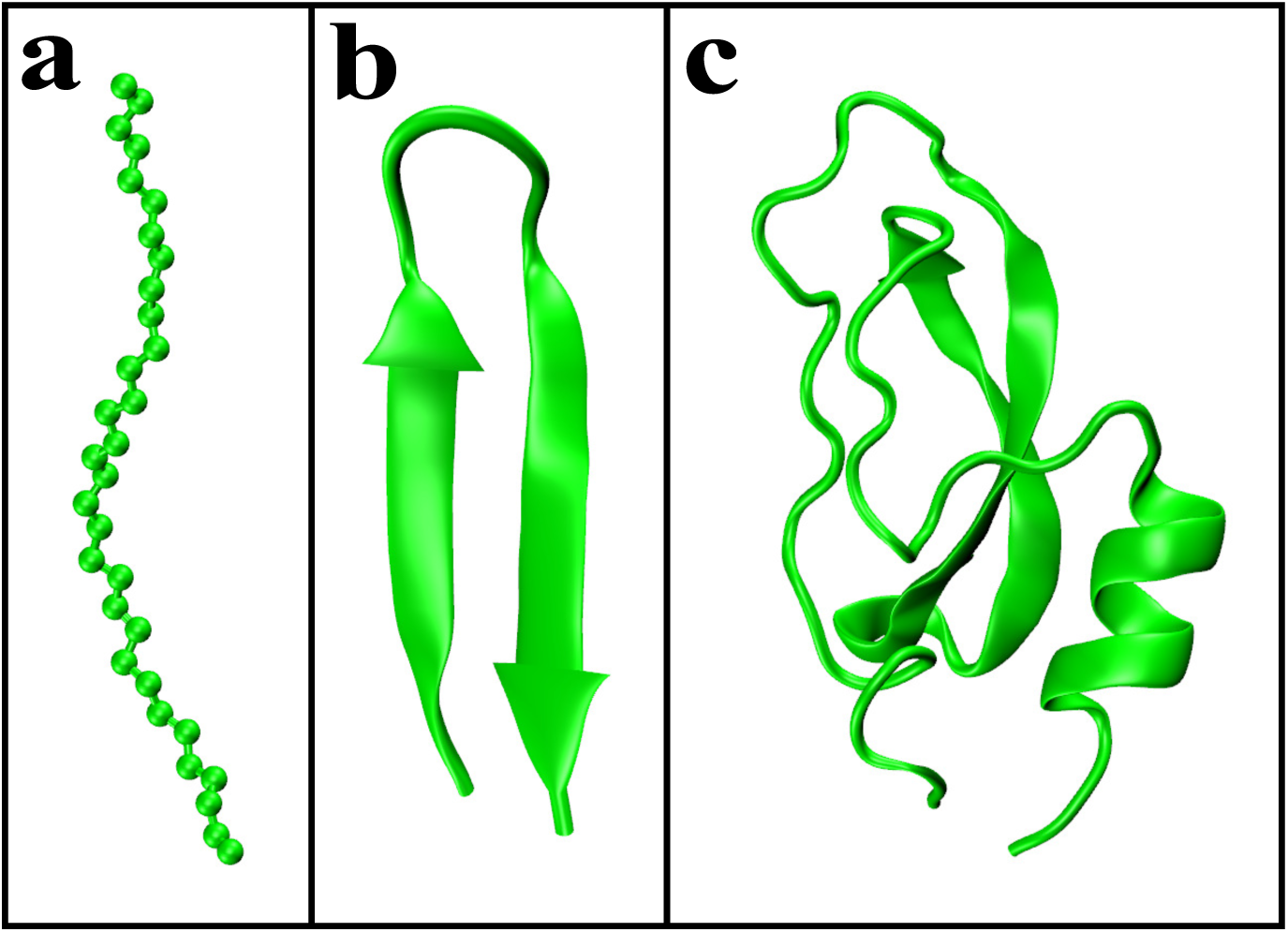
Computational models of the three systems studied in the current work: a) 32-bead model polymer, b) C-terminal domain of GB1 *β*-hairpin (residues 41–56) and c)Bovine pancreatic trypsin inhibitor (BPTI) protein

The 16-residue long peptide Ace-^41^GEWTYDDATKTFTVTE^56^-Nme, as taken out from the full-length GB1 protein NMR structure (PDB: 1GB1^31^) was modelled using AMBER03-STAR^32,33^ forcefield and solvated in a truncated octahedron simulation cell containing 984 TIP3P^34^ water molecules and electroneutralized. The current work utilizes our previously run 200 short MD trajectories of the *β*-hairpin, each 100 ns long and spawned from prior replica exchange molecular dynamics (REMD) derived trajectories.^7^ In addition, we have spawned 200 more MD trajectories (and hence a total of 400 trajectories) to improve the statistics of constructed conformational landscape for *β*-hairpin. The readers are referred to work of Mondal and coworkers^7^ for simulation details.

The simulation trajectories for BPTI correspond to prior work of Shaw and coworkers ^35^ and were generously provided by D. E. Shaw research.

All frameworks of Time structured Independent Component Analysis (TICA) and Markov state model (MSM) were developed using pyemma. ^36^

## Results and Discussion

This communication illustrates that GNAP encodes relevant information concerning conformational folding of a macromolecule and in combination with TICA, can serve as an efficient collective variable. Towards this end, we first consider a model 32-bead charge-neutral polymer with equal number of beads having partial charges of opposite polarity (see representative snapshot in Figure 1c and Figure 2 a)). Recent MD simulation by Mondal and coworkers^29^ (also see simulation methods) has shown that this polymer can undergo reversible collapsed⇌extended transition in aqueous media and the transition often involves a hairpin-like intermediate as well. We calculate the time profile of GNAP of the polymer from its representative MD simulation trajectories. As depicted in Figure 1c, we find that GNAP captures the dynamic evolution of the polymer with time and responds in unison with conformational fluctuation of the polymer. Specifically, we find that, starting with collapsed conformation as reference, the value of GNAP increases with transition from collapsed to hairpin or extended conformation and vice versa (Figure 1d)

The observation that GNAP of the polymer can dynamically evolve, in sync with its conformational fluctuation, encouraged us to explore the power of GNAP in mapping the free energy landscape of a macromolecule. However, one of the key features of GNAP is that its computation depends on a reference structure. For a conformationally floppy macromolecule, the choice of reference structure may not always be unique, which in turn influences the reconstruction of the free energy landscape. This is demonstrated by the supplementary Figure S1. To circumvent the dependence of GNAP on single reference structure, we here develop a robust protocol by combining techniques of TICA^24,25^ with GNAP. Our proposed work flow is enumerated in Figure 3. The main purpose of this protocol is to optimally combine the GNAPs obtained from multiple reference configurations. Towards this end, we first perform adaptive molecular dynamics sampling of the macromolecule of interest. The aggregated conformations resulting from the adaptive sampling is subsequently subjected to K-mean clustering. ^37–39^ The cluster centres thereby obtained provide us with a set of representative reference structures, each of which is used to compute individual GNAP. These multiple GNAP values, corresponding to as many reference structures, are then optimally combined in a linear fashion within the frame-work of TICA. As has been shown earlier by Pande and Noe, ^24,25^ TICA efficiently optimises the coefficients of the linear combination by maximising the auto-correlation of projection. The number of TICA dimensions gets further reduced by choosing only those which conserves 90% of the kinetic-variance.

**Figure 3:**
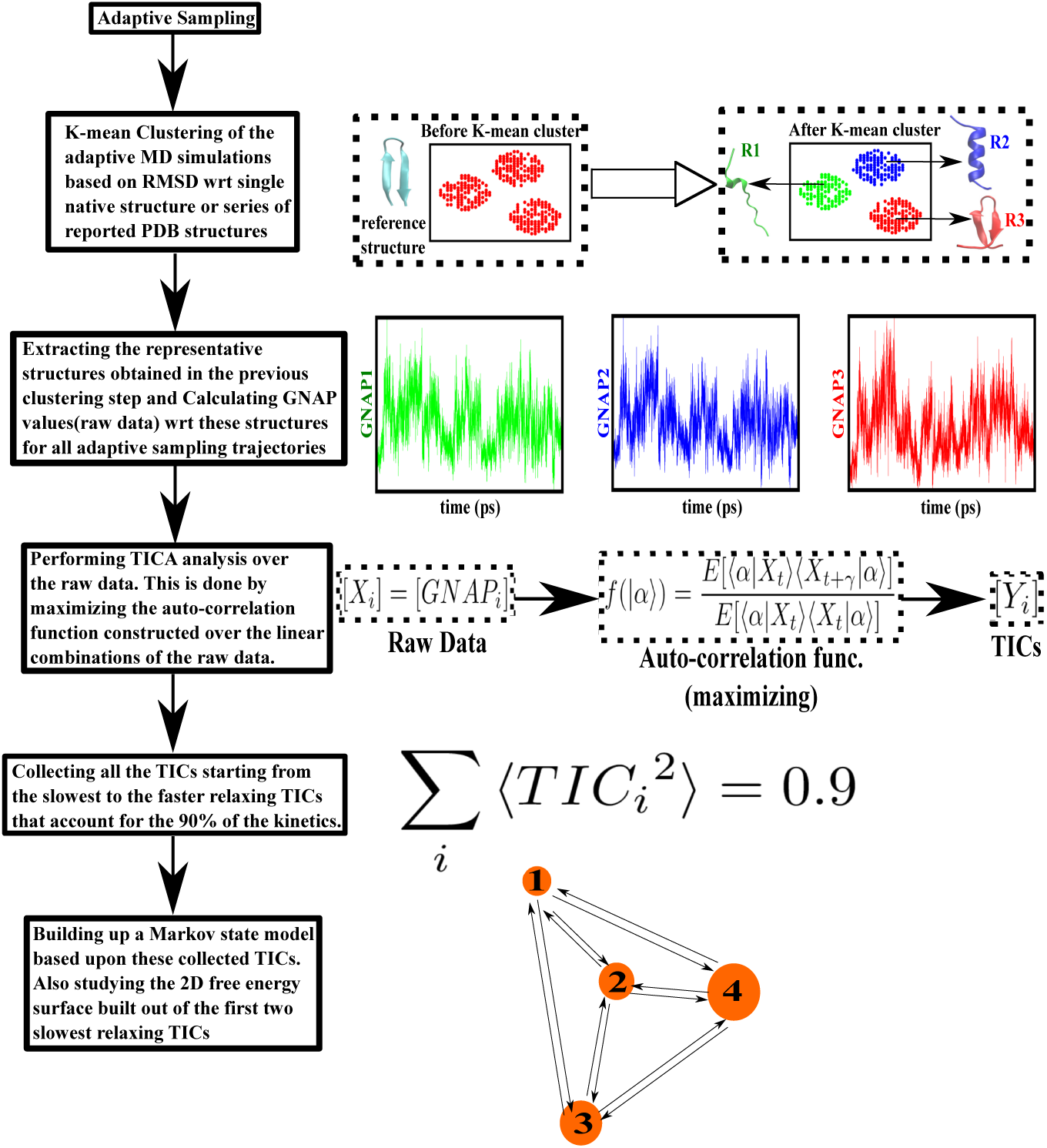
Scheme representing the computational protocol described in the current work. The trajectories based on adapting sampling is the source of the data points. The data points are first subjected to K-mean clustering. The GNAP calculation requires a reference structure. The output of the K-mean clustering i.e. the representative structures of the clusters are used as the different structures and the GNAP values so obtained act as the raw data for the TICA step. Collecting such TICA components which account for the 90 percent of the kinetics of the system is required to properly build a markov state model.

When this strategy was applied for the model polymer discussed earlier, a set of 12 reference structures were found to exhaustively cluster the conformational ensembles and hence with these 12 reference structures we calculate a set 12 GNAP values for our adaptive sampling trajectories. In the case of model polymer, we noted that just 3 out of 12 TICA dimensions were sufficient to conserve 90% of the kinetic-variance. Figure 4a projects the two-dimensional free energy landscape of the model polymer along top two slowest linear combinations of GNAPs, namely *TIC*_1_ and *TIC*_2_. The free energy surface is obtained after reweighing the probability distribution by the stationary distributions. We find that the optimal linear combinations of GNAP are able to fully resolve the landscape of the model polymer into multiple key conformations (figure 4c), namely extended (state-0, 34.39%), collapsed (state-1, 23.29%), distorted hair-pin (state-2, 7.16%) and symmetric hair pin (state-3, 35.13%), which comes out to be most populated conformation. Reconstructed transition networks (figure 4b) among these GNAP-projected key polymer conformations, as obtained via a Markov state model (MSM) analysis^40^ (over 800 discrete micro states at 250 ps lag time), show diverse array of rapid state-to-state transitions, especially between collapsed and perfect hairpin structure.

**Figure 4:**
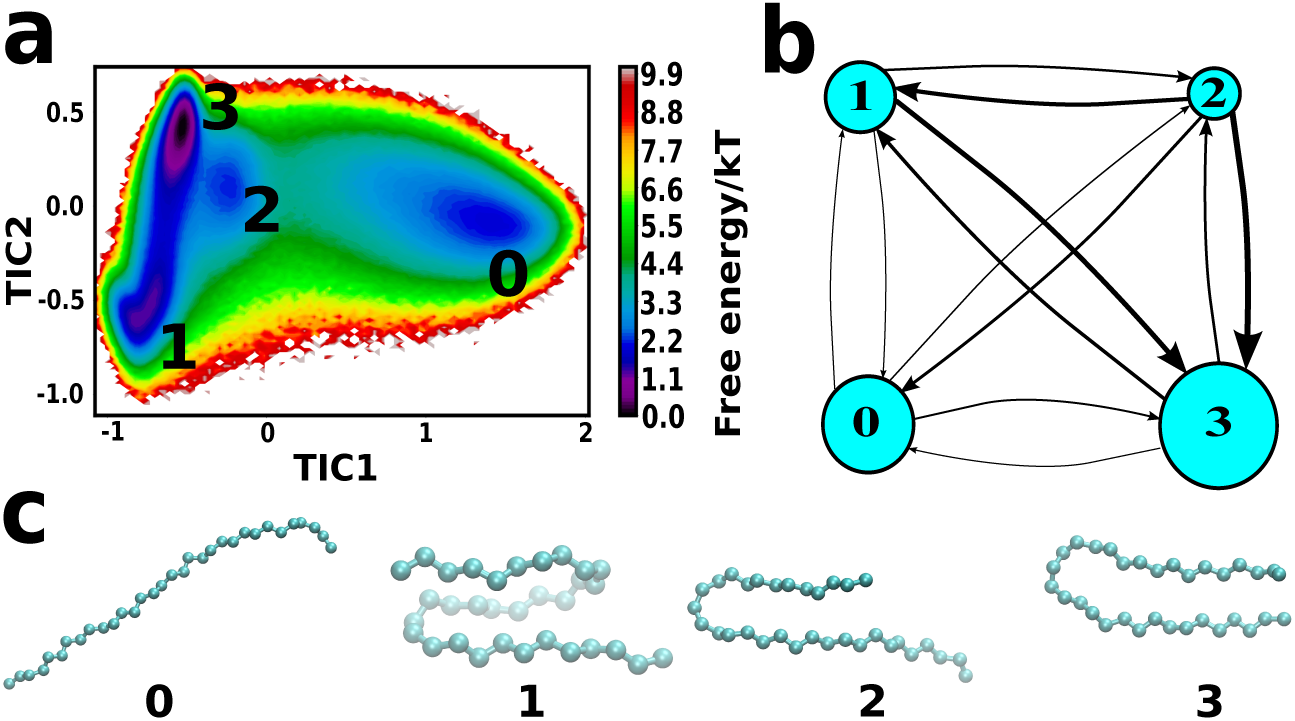
The 4 state markov model for the 32 bead polymer chain. **a**. Free energy profile along the two underlying slowest mode of relaxation reveal four minima. **b** the various fluxes connecting the four minima, with the thickness of the arrows representing the relative fluxes. **c**. the conformational macrostate at the four minima. The hairpin structure is the most populous and the collapsed structure is the next most populous structure. Also there a state which is a distorted version of the hairpin very close by the perfect hairpin structure.

The ability of GNAP in exhaustive quantification of underlying conformational landscape becomes more prominent in hierarchically more complex systems than the model polymer : *β*-hairpin GB1 peptide and Bovine pancreatic trypsin inhibitor (BPTI) protein (see Figure 2b and 2c). In particular, for GB1 peptide,(see simulation methods and our earlier work ^7^ for details of simulation models and methods), we find that (see Figure 5) the combination of GNAP together with the protocol described in Figure 3 brings out the complex free energy landscape along the slowest and second slowest dimensions. Subsequent MSM analysis (based on 1000 discrete micro states at a lag time of 600 ps demonstrates that a 5-state model with state-0 being the symmetrically distorted beta hair-pin (14.85%), state-1, perfect beta hairpin (49.85%, state-2: assymetrically distorted beta hair-pin (7.10%), state-3: terminal *α*-helix (3.67%) and state-4: central *α*-helix (24.51%) aptly describes the overall landscape (see figure 5). The transition-network shows that flux from state 1 to any other state, especially state 1-to-3 are extremely feeble, implying that the transformation from beta hair-pin to terminal helix is most rare and difficult. This is also backed up by our common understanding that such a transformation requires a huge structural re-arrangement at multiple sites.^7^

**Figure 5:**
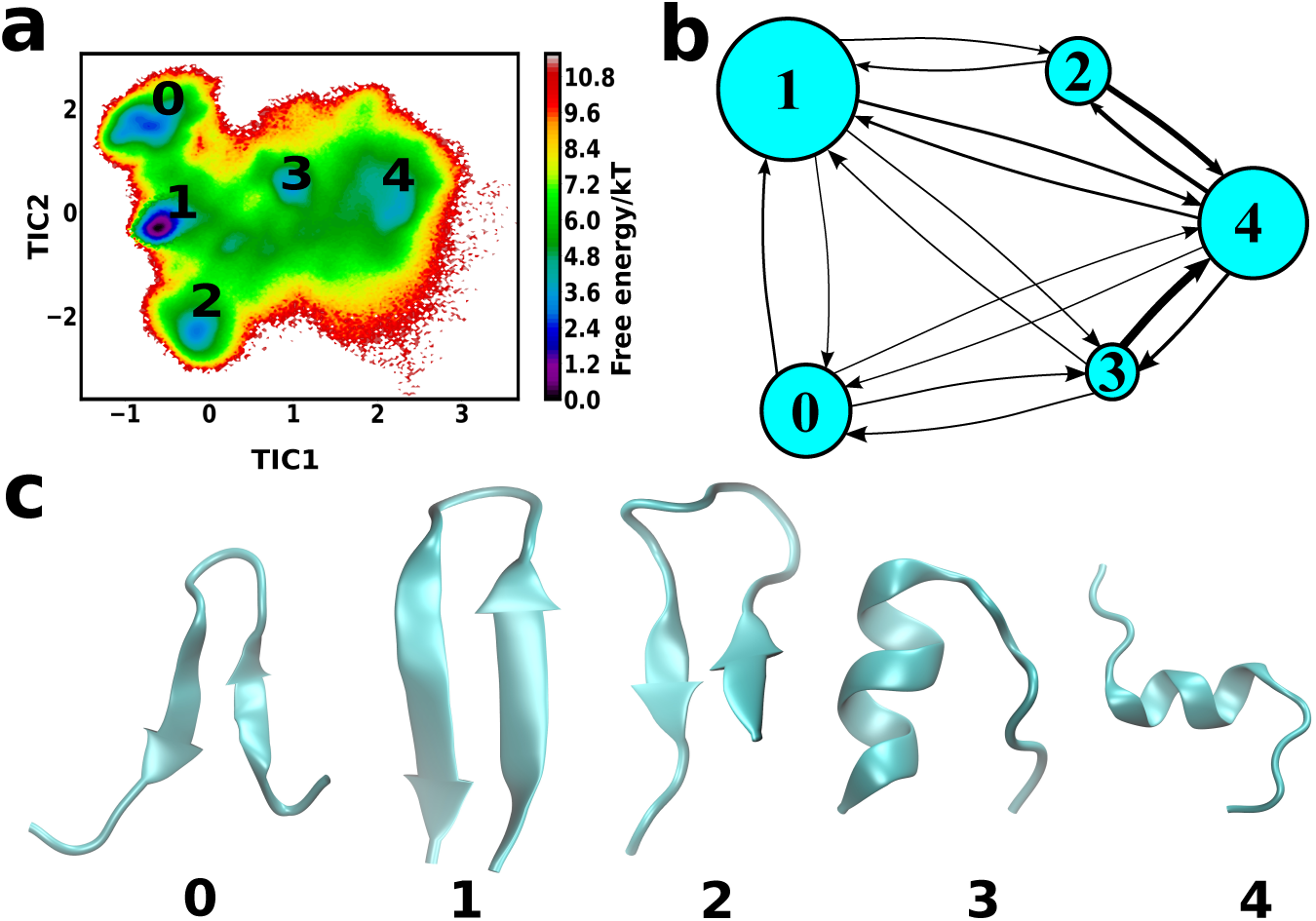
**a.** The 5 state markov model for the GB1 protein, mapped on the free energy surface along TIC1 and TIC2 (see text for details of TIC1 and TIC2) **b**. Transition network among the macro-states (the thickness of the arrows represents the relative fluxes). The free-energy profile recovers that the transitions among the structurally similar conformations 0, 1 and 2 (these structures lie along the second slowest modes) are somewhat faster compared to those which are structurally quite distinct which requires huge structural transformation **c**. Snapshot of structures corresponding to the minima.

An important aspect of the current formalism is that we can investigate the most probable non-affine modes by projecting out homogenous fluctuations from the local displacements. These non-affine modes can potentially infer local structural changes between a pair of macro states. We illustrate this via the supplemental movie S1 which demonstrates the non-affine modes responsible for transition of GB1 from beta hair-pin structure to the central helix structure. The movie also demonstrates a key advantage of GNAP over popular approach of principal component analysis (PCA) (see supplemental movie S2 for PCA):^41,42^ the current approach can explore local changes and the projection on the non-affine subspace ensures that all trivial motions are filtered out and thereby it is possible to retrieve and analyse predominant modes responsible for the conformational changes.Figure 6 quantifies the key difference between PCA and GNAP-analysis for the above-mentioned sampling of GB1-hairpin-transition from *β*-sheet to central *α*-helix. In Figure 6a, the spatial distribution of the normalized NAP-susceptibility^16,23^ across the protein molecule is calculated over the aforementioned sampling. The two particles in red and also circled in red are the centers of high NAP-susceptibility and therefore likely to produce non-afffine conformational changes. To explore the difference between PCA and NAP on equal footing, we solve the eigenvalue equation of their individual covariance matrices. The eigen spectra at the two locations, as obtained by current formalism, are shown in Figure 6b where it is clearly seen that the highest eigenmode is completely separated from the rest of the eigen-spectrum for the two high NAP-susceptibility regions. This large separation confirms that the modes associated with the separated eigenvalues are indeed non-random. On the other hand, the PCA-derived eigen spectrum (Figure 6b) shows a relatively lower gap in its associated eigenvalues than those obtained in case of NAP-derived covariance matrix. A comparison of displacement modes at these two locations (Figure 6c) derived from NAP analysis and from PCA analysis shows that the NAP analysis recovers the helical curvatures and the twists that are required to attain the central *α*-helix begin to appear, while the PCA mode provides a much limited resolution in regard with local mechanism involved in the transition from hair-pin to central alpha helix. It can also be seen from Figure 6d that the net effective displacement vector involved in the transition from *β*-sheet to central *α*-helix conformation projected over the most prominent eigenmodes (indicated by the highest eigen values) in the two methods is higher in the case of NAP-analysis than PCA-analysis, which suggests that the modes obtained from the NAP-analysis are able to mimic the underlying dynamics involved in this transition more closely than the PCA analysis.

**Figure 6:**
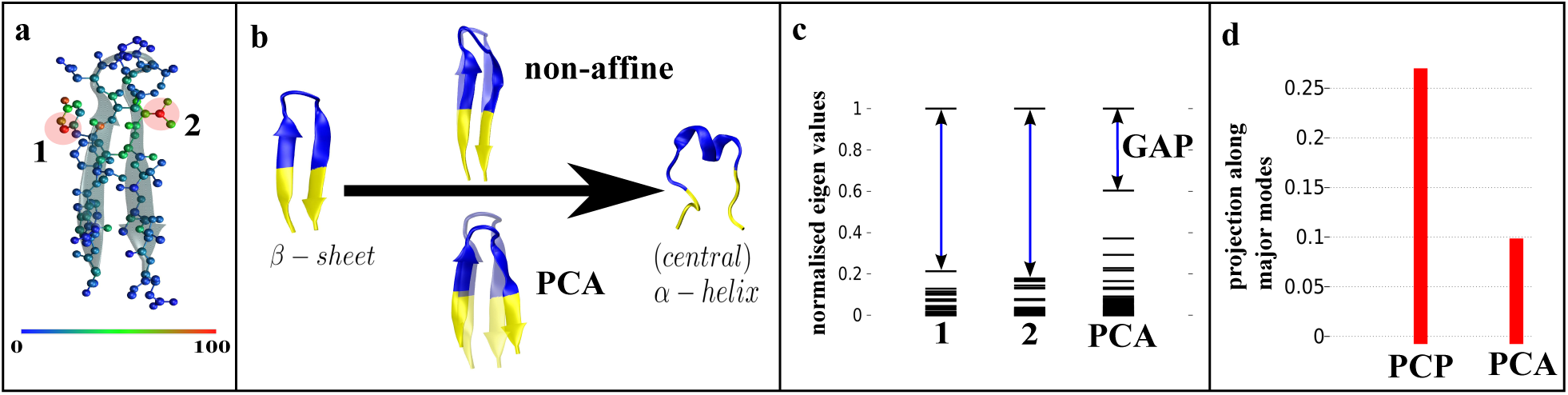
Comparison between GNAP and PCA analysis for GB1 protein system **a**. Spatial distribution of NAP susceptibility. This analysis provides the hotspots region at indicated regions in protein (the regions, locations 1 and 2, that are red in colour). **b**. The non-affine displacement modes derived from the GNAP analysis better manifest the helical topologies required for going from *β*-sheet to central *α*-helix conformation than PCA. **c**. Comparison of eigen-spectrum obtained from NAP analysis at the two locations 1 and 2 with the PCA eigen-spectrum for the entire GB1 system. **d**. The quantification of how well NAP analysis predicts the transition process, compared to the PCA analysis, via the projection of the Δ_*transition*_ (defined as the difference between the coordinates of the *β*-sheet and central *α*-helix conformations) on the highest eigen value modes obtained in the two methods.

Finally, going beyond simple polymers and peptides, GNAP is found to aptly describe the conformational phase-space for prototypical protein BPTI (Figure 7). The milli-second long simulations by D.E. Shaw group^35^ have earlier provided atomic-scale insights into the folding/unfolding events of BPTI. The projection of the trajectory (generously provided by D. E. Shaw research) along the top two TICA-derived slowest GNAP-combination reveals multiple key intermediates of BPTI (Figure 7a). Interestingly, similar projection along more typical CV namely RMSD fails to resolve all the key intermediates (figure 7b) The macro state corresponding to folded native conformation automatically appears as the free energetically most stable state (state 1) in the GNAP reconstructed free energy projection. At this minima, we recover a structure with left-handed conformation of the disulfide bridge between Cys14 and Cys38. Interestingly, this has been previously reported by Shaw and coworkers as the most populated state. On the other hand, state 2 represents conformation with large deviation from native structure, which is the unfolded conformation. Most importantly, the GNAP analysis recovers the well-known “left-handed” conformation of the disulfide bridge between Cys14 and Cys38 in state 1, 2 and 4, as earlier reported by simulations and NMR experiments. Finally, state 3 brings out the ‘right handed’ conformation ^43,44^ (see Grey et al^43^ and captions of Figure 7 for details.)

**Figure 7:**
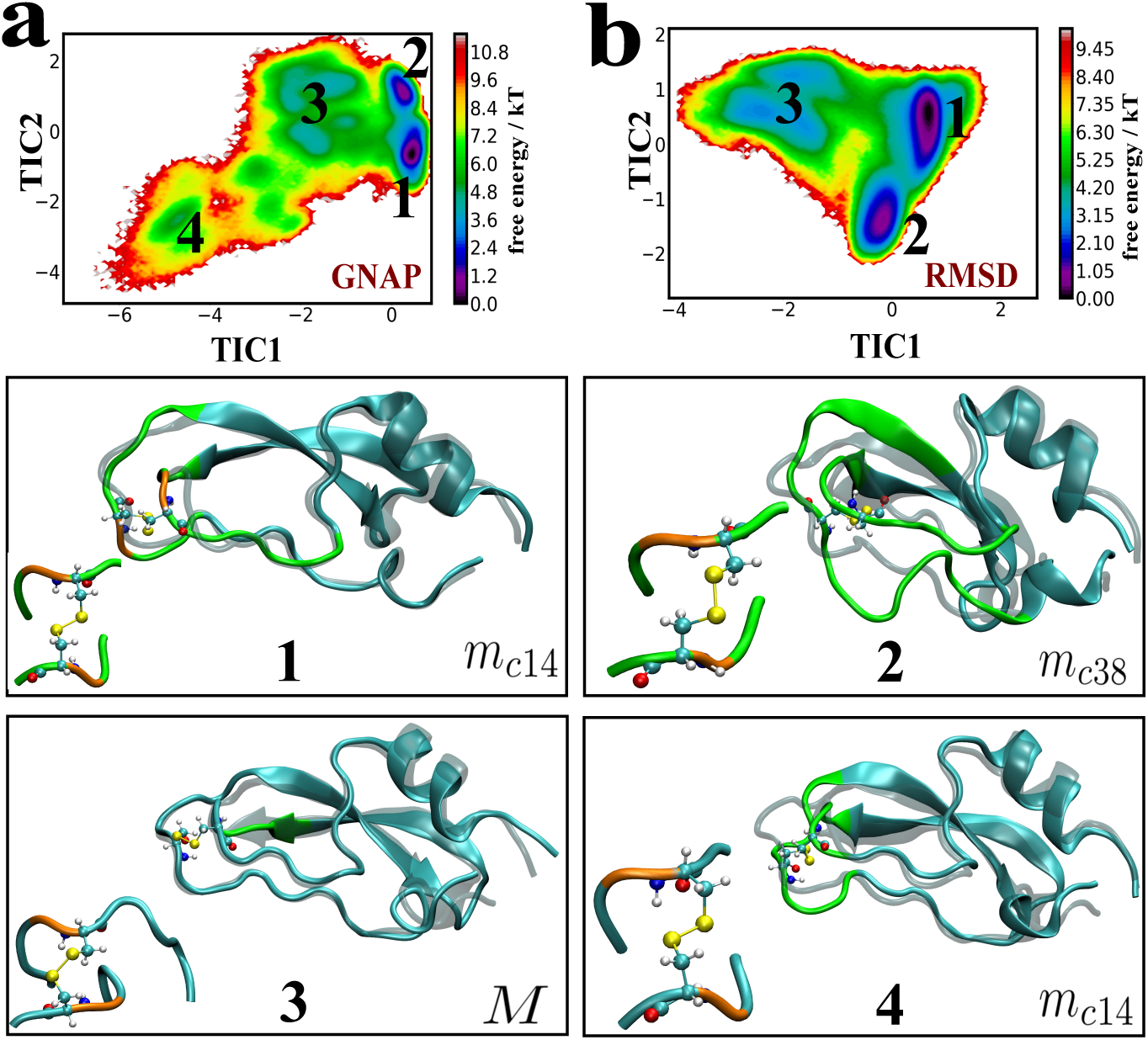
Comparison of GNAP with RMSD for BPTI system **a**. The free-energy profile along TIC1 (slowest mode of relaxation) and TIC2 (the second slowest mode of relaxation) constructed out of linear combination of multiple GNAPs corresponding to representative reference structures obtained in the K-mean cluster step **b**. The free-energy profile along TIC1 and TIC2 constructed out of linear combination of RMSDs relative to the same sets of reference representative structures, as used in case of GNAPs. The four structures 1, 2, 3 and 4 have the same reference structures(in grey ghost representation) while the structure from the four minima are shown in cyan with distorted part in green. M (structure 3) is the right-handed disulphide conformation and *m*_*c*14_ (structure 1 and 4 (with distortion)) and *m*_*c*38_ (structure 2) are the left-handed disulphide conformation, as reported by Grey et al.^43^ Structure 4 is not captured in the RMSD-TICA free-energy profile.

## Conclusion

In summary, the current work demonstrates that the non-affine displacements in bio macro-molecule can act as a good marker for characterising its key conformational fluctuations. Specifically, we formulate a global non-affine parameter (GNAP) by averaging over all local non-affine displacements corresponding to each protein heavy atoms and we show that GNAP in turn can capture the essential collective motion in bio-macromolecules. We also find that GNAP, when used in combination with TICA, judiciously handles the choice of multiple reference states and resolves the underlying free energy landscape into key basins. These are efficiently demonstrated for a series macromolecules of hierarchical complexity (model polymer, peptides and proteins). More over, GNAP is found to provide better resolution of the free energy landscape of a protein BPTI compared to RMSD, a conventional CV regularly used for projecting the biomacromolecular motion. The non-affine modes, obtained by projecting out homogenous fluctuations from the local displacements, are found to be responsible for local structural changes required for transitioning between a pair of macro states. Finally, we show that a local NAP analysis can be used to filter out the essential macromolecular configurations responsible for slowly evolving conformational changes such as that from a *β*-sheet to *α*-helix, pinpointing the residues where such a change is likely to initiate. Overall, identification of non-affine fluctuations in protein offers a distinct advantage for understanding the bio-macromolecular dynamics and its quantifier GNAP holds the promise of acting as an optimal CV for projecting the free energy landscape. The availability of an analytical expression for GNAP makes it possible to patch it with any standard MD analysis tool for post-simulation analysis. Along this line, the current work has seen the implementation of GNAP into popular open access MD analysis tool PLUMED.^27^ This holds future promises of driving a biophysical process along its collective non-affine displacement using enhanced sampling techniques.

## Acknowledgement

This work was supported by computing resources obtained from shared facility of TIFR Center for Interdisciplinary Sciences, India. JM would like to acknowledge research intramural research grants obtained from TIFR, India, Ramanujan Fellowship and Early Career Research funds provided by the Department of Science and Technology (DST) of India (ECR/2016/000672). We thank D. E. Shaw research for generous access to their simulation trajectories of Bovine pancreatic trypsin inhibitor.

## Supporting Information (SI)

Figure demonstrating the variation of free energy due to different choice of reference structures(PDF). Movies showing non-affine modes and PCA modes for transition from beta-sheet to central helix of GB1 peptide.(mpeg)

## TOC Graphic

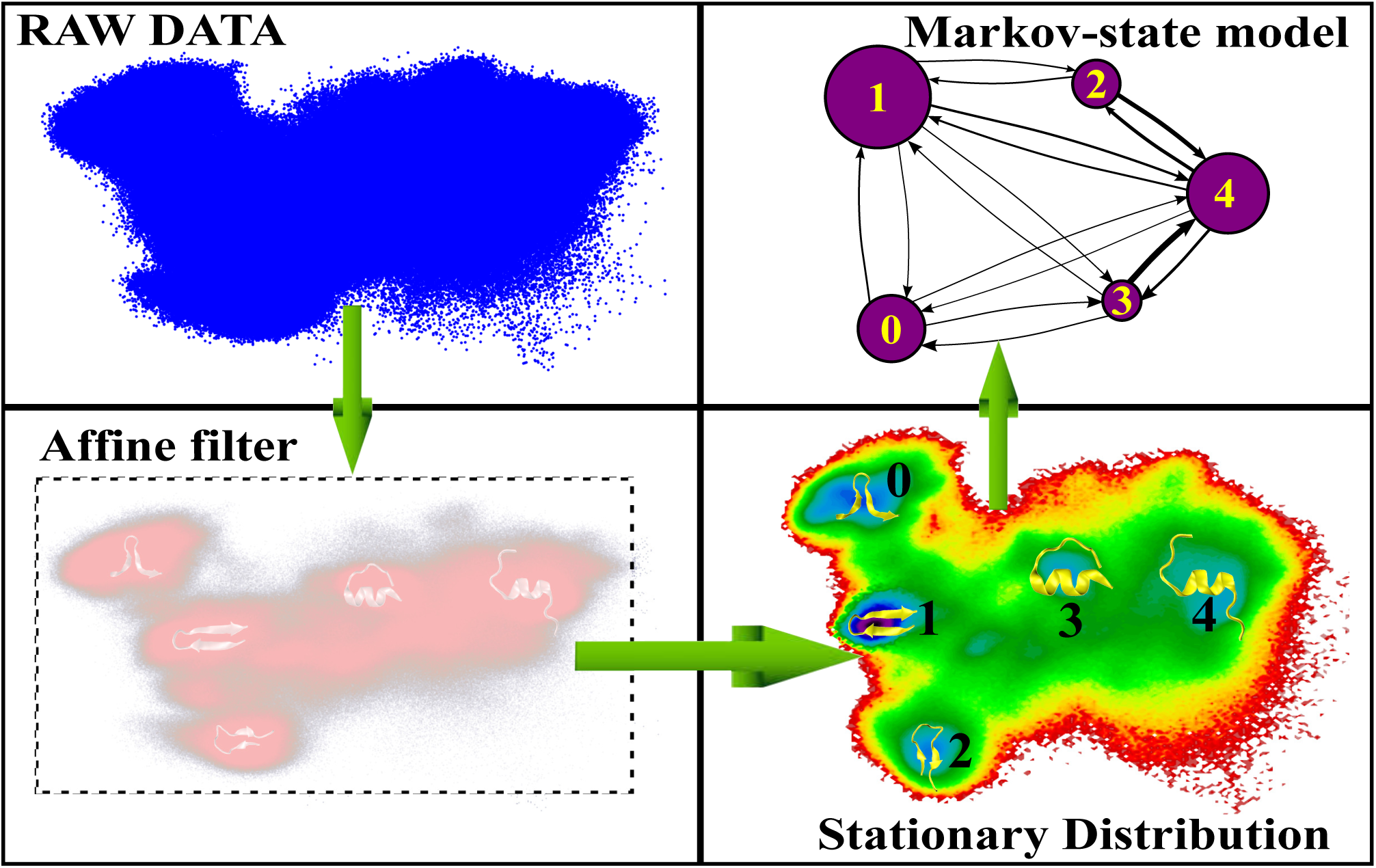

