## Supplementary figures for "Non-affine displacements encode collective conformational fluctuations in proteins"

### Supporting Information for “Non-affine displacements encode collective conformational fluctuations in proteins”

#### Description of Supplemental Movies:

- Movie S1 demonstrates the non-affine modes responsible for transition of GB1 from beta hair-pin structure to the central helix structure.

- Movie S2 demonstrates the PCA models responsible for transition of GB1 from beta hair-pin structure to the central helix structure.

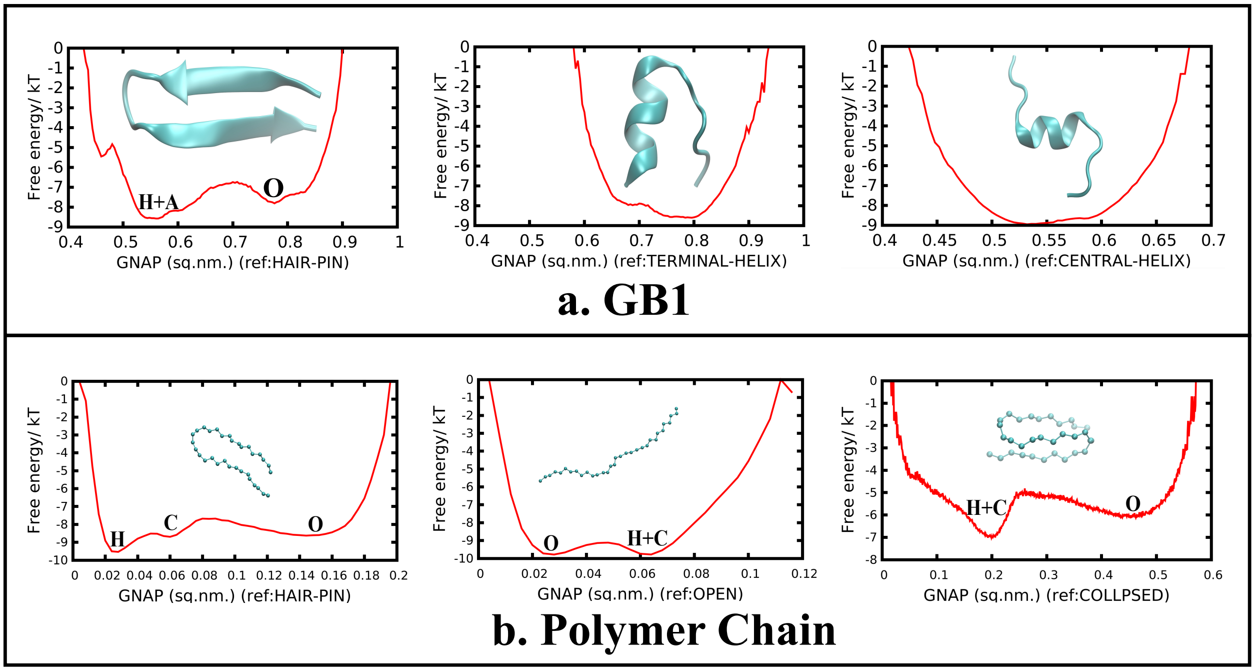


Figure S1: Illustration of different free energy curve arising due to choice of different reference structure for computing GNAP.
